# Identifying patterns differing between high-dimensional datasets with generalized contrastive PCA

**DOI:** 10.1101/2024.08.08.607264

**Authors:** Eliezyer Fermino de Oliveira, Pranjal Garg, Jens Hjerling-Leffler, Renata Batista-Brito, Lucas Sjulson

## Abstract

High-dimensional data have become ubiquitous in the biological sciences, and it is often desirable to compare two datasets collected under different experimental conditions to extract low-dimensional patterns enriched in one condition. However, traditional dimensionality reduction techniques cannot accomplish this because they operate on only one dataset. Contrastive principal component analysis (cPCA) has been proposed to address this problem, but it has seen little adoption because it requires tuning a hyperparameter resulting in multiple solutions, with no way of knowing which is correct. Moreover, cPCA uses foreground and background conditions that are treated differently, making it ill-suited to compare two experimental conditions symmetrically. Here we describe the development of generalized contrastive PCA (gcPCA), a flexible hyperparameter-free approach that solves these problems. We first provide analyses explaining why cPCA requires a hyperparameter and how gcPCA avoids this requirement. We then describe an open-source gcPCA toolbox containing Python and MATLAB implementations of several variants of gcPCA tailored for different scenarios. Finally, we demonstrate the utility of gcPCA in analyzing diverse high-dimensional biological data, revealing unsupervised detection of hippocampal replay in neurophysiological recordings and heterogeneity of type II diabetes in single-cell RNA sequencing data. As a fast, robust, and easy-to-use comparison method, gcPCA provides a valuable resource facilitating the analysis of diverse high-dimensional datasets to gain new insights into complex biological phenomena.

## Introduction

Investigators in the biological sciences are increasingly collecting high-dimensional datasets that are challenging to analyze, with modalities ranging from imaging to electrophysiology to single-cell RNA sequencing. Dimensionality reduction algorithms such as principal components analysis (PCA) and its many variants (***Pearson, 1901; Hotelling, 1933; Zou et al., 2006; Zass and Shashua, 2006; Tipping and Bishop, 1999***) are used widely to help simplify these datasets and facilitate analysis. PCA examines the covariance structure of the data to find dimensions that account for more variance than chance; these constitute patterns that are overrepresented in the data, such as assemblies of neurons whose activity fluctuates up and down together across time in a neural recording (***Chapin and Nicolelis, 1999; Peyrache et al., 2010; Lopes-dos Santos et al., 2013; Sjulson et al., 2018***), or networks of genes that are up- or down-regulated together across cells in a single-cell RNAseq dataset (***Chung and Storey, 2015; Li et al., 2016***). However, in many cases, the goal is to compare data collected under two different experimental conditions, which we refer to here as datasets. Since PCA and other dimensionality reduction techniques operate on only one dataset, they cannot take experimental conditions into account.

The most common approach for comparing two high-dimensional datasets is linear discriminant analysis (***Izenman, 2008***) or its multidimensional analog, partial least squares discriminant analysis (***Brereton and Lloyd, 2014***). These methods find dimensions that optimally distinguish one dataset from the other, which could correspond to which neurons fire more, or which genes are upregulated, in condition *A* vs. condition *B*. However, an analogous method to compare the covariance structure of two datasets is not as well established. This addresses more subtle and detailed questions, such as which subsets of neurons exhibit increased temporal correlations in condition *A* than *B*, or which subsets of genes are more likely to be up- or downregulated together in individual cells in condition *A* than *B*. Mathematically, answering these questions corresponds to finding dimensions that account for more or less variance in *A* than *B*.

Recently, contrastive PCA (cPCA) was proposed as a method to address this problem (***Abid et al., 2018***). Although cPCA is an important first step, it requires a hyperparameter *α*, which controls how much covariance from the second condition to subtract from the first. The algorithm must therefore iterate over multiple choices of *α* with no objective criteria to determine which value of *α* yields the correct answer. Moreover, cPCA is asymmetric, identifying the most enriched dimensions in the first condition after subtracting out the second condition as background; it cannot treat the two experimental conditions equally.

Here we propose a novel solution to these problems we call generalized contrastive PCA (gcPCA). We first demonstrate the role the *α* hyperparameter plays in cPCA, then explain our strategy for eliminating it. We then describe an open source toolbox for Python and MATLAB implementing several versions of gcPCA with different objective functions that are either asymmetric or symmetric, orthogonal or non-orthogonal, or sparse or dense, tailored to suit the specific application at hand. Finally, we demonstrate the utility of gcPCA in the analysis of diverse biological datasets.

## Results

### The cPCA hyperparameter *α* compensates for bias toward high-variance dimensions in noisy, finitely-sampled datasets

To explain the need for the hyperparameter *α* in cPCA and how we avoid it in gcPCA, we will describe the objective function of each method and show how they perform in generated synthetic data. For illustration purposes, we generated synthetic data for two experimental conditions containing two-dimensional manifolds on a background of high-variance shared dimensions. The generated data consisted of condition *A*, with a manifold (additional variance) in the 71st and 72nd dimensions (ranked in order of descending variance), and condition *B*, with a manifold in the 81st and 82nd dimensions (Fig. 1A). The manifold dimensions contained less total variance than most of the other dimensions in the dataset, but their variance is two-fold higher in one condition relative to the other (i.e., the 81st and 82nd dimensions have twice as much variance in condition *B* than condition *A*). An important property of real-world biological datasets is that they are noisy and finitely sampled. We aimed to model the finite data regime by comparing 1 × 10^3^ samples (finite data) to 1 × 10^5^ samples, which approximates infinite data. To inspect the effects of finite sampling on estimated variance, we projected the “finite” and “infinite” data onto the ground truth dimensions and calculated the variance explained by each (see methods). Our results (Fig. 1B) reveal that finite sampling yields noisy estimates of the true variance, with greater noise in high-variance dimensions.

**Figure 1.**
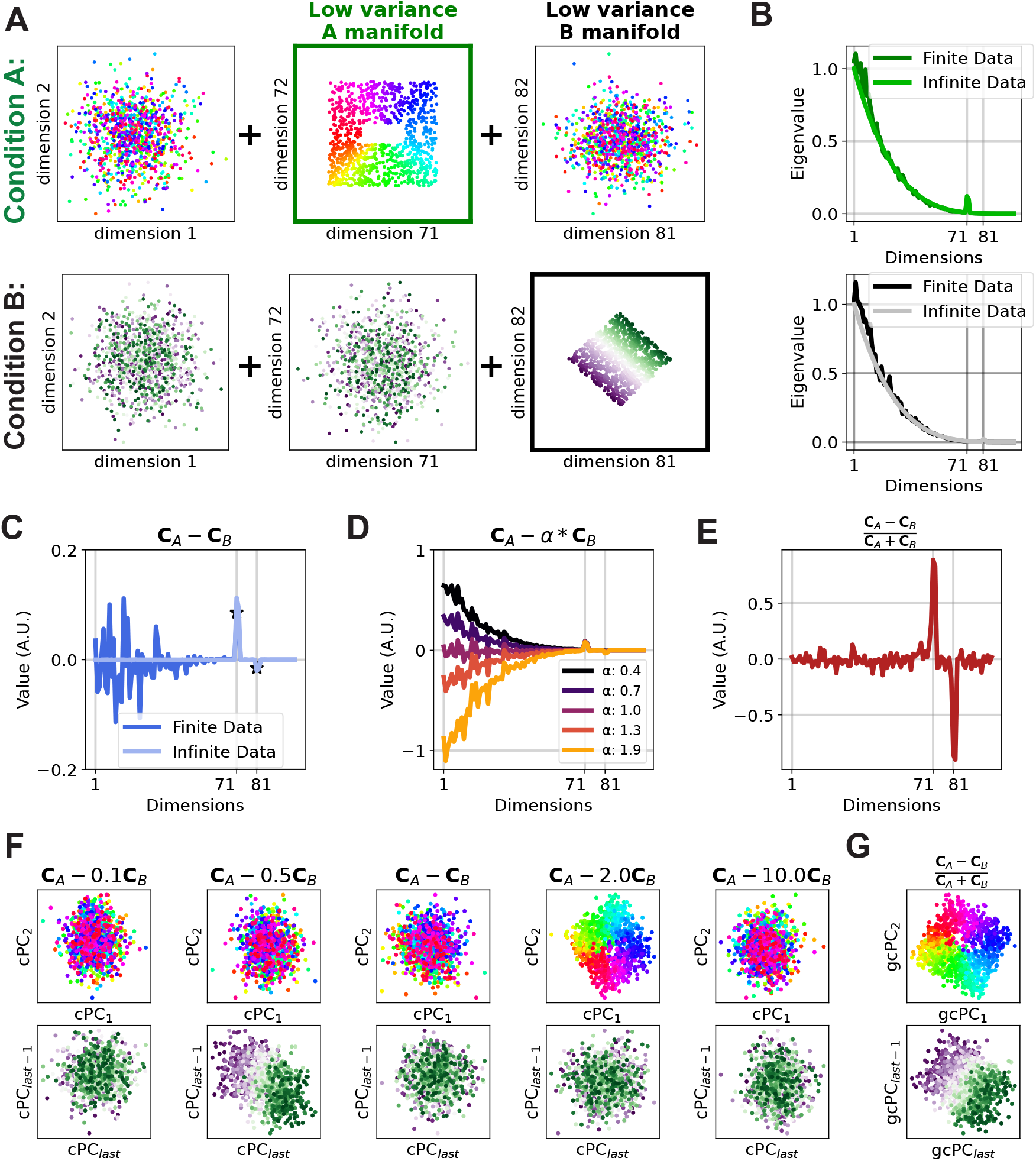
Generalized contrastive PCA avoids cPCA’s need for the hyperparameter *α* in noisy, finitely-sampled data. **A** We generated two conditions in noisy synthetic data that each contain low-variance manifolds that are not present in the other. These manifolds have lower overall variance than many other dimensions and are not trivially discoverable. **B** Eigenvalue spectra for each condition estimated from finite (dark line) or infinite (light line) data. Note the sampling error in the finite date case. **C** With infinite data, eigendecomposition of (**C**_*A*_ − **C**_*B*_) suffices to extract the correct answers (dimensions 71-72 and 81-82, light lines). However, with finite data, these peaks are smaller than the sampling error in high-variance dimensions, creating a bias toward high-variance dimensions being selected. **D** cPCA uses the hyperparameter *α* to adjust how much influence **C**_*B*_ has on **C**_*A*_. As *α* increases, the bias toward high-variance dimensions decreases until it becomes negative with *α* > 1, eventually exposing the differences in lower-variance dimensions. Importantly, there is no way to know which value of *α* yields the correct solution. **E** Using gcPCA, the dimensions most changed in each condition are identified correctly, even with finitely-sampled data. **F** cPCA with the optimal choice of *α* does not extract the correct dimensions in *B*. **G** gcPCA identifies the enriched dimensions in each condition and correctly returns the low-variance manifolds. Because gcPCA is symmetric, it extracts the correct dimensions in both *A* and *B*.

To understand the practical consequences of this, it is helpful to start by reviewing traditional PCA. With PCA, the principal components of a data matrix **D**, of size *n* × *p* (samples × features), are the dimensions explaining the most variance. These can be identified by estimating the covariance matrix **C** = **D**^⊺^**D**/(*n* − 1), then solving the following quadratic optimization problem in equation 1:

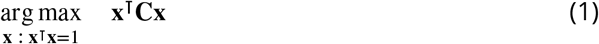

This problem can be solved by eigendecomposition of **C**, yielding the matrix of eigenvectors **X** known as principal components (PCs).

To extend this to two datasets, a logical strategy involves formulating an objective function to describe the difference in variances between the two conditions, enabling us to extract dimensions that show the greatest increase in variance in *A* relative to *B*. We now have two covariance matrices, **C**_*A*_ and **C**_*B*_, and the contrastive PCs (cPCs) are the vectors that maximize the objective function 2:

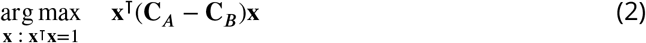

Analogous to traditional PCA, this problem can be solved by eigendecomposition of (**C**_*A*_ − **C**_*B*_). This yields cPCs that account for more variance in either condition *A* or *B*, corresponding to the eigenvectors with the largest or smallest eigenvalues, respectively. With infinite data, this approach correctly finds the dimensions enriched in conditions *A* and *B* (Fig. 1C, light line), but with finite data, it fails to do so because the sampling error in higher-variance dimensions is larger than the true signal in the lower-variance dimensions (Fig. 1C, dark line). In other words, it has a systematic bias toward high-variance dimensions. To compensate for this effect, cPCA ***Abid et al. (2018)*** introduces the hyperparameter *α*, changing the following objective function to 3:

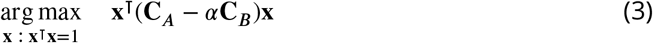

As discussed in ***Abid et al. (2018)***, *α* represents a trade-off determining the extent to which **C**_*B*_ influences the identification of enriched vectors in **C**_*A*_. In our synthetic data, we can visually appreciate the effect of different values of *α* in the resulting cPCA objective function value (Fig. 1D). In effect, *α* tunes the amount of bias toward high-variance dimensions in the cPCA calculation, with *α* < 1 biasing toward high-variance dimensions and *α* > 1 biasing against them. In this case, *α* = 2 yields the correct solution that dimensions 71-72 are enriched in *A* (Fig. 1F), but other values of *α* yield equally plausible, but incorrect, solutions. Importantly, we can only determine which solution is correct because we knew the answer in advance, which is not typically the case for experimental data. Further, negative values in cPCA are generally interpreted as dimensions enriched in condition *B* (***Abid et al., 2018; Boileau et al., 2020***), but our simulation shows that values of *α* larger than 1 bias the highest-variance dimensions to be negative (Fig. 1D). This creates the illusion that these dimensions are enriched in condition *B*, even though the correct answer is that only dimensions 81-82 are enriched in *B*. Similar to the situation of finding dimensions enriched in *A*, the results depend on the choice of *α*, with no way to determine which solution is correct (Fig. 1F). Importantly, the range of *α*’s yielding the correct solution can be incredibly narrow, as in Fig. S1, where *α* = 2.6 yields the correct solution, but 2.2 or 3.0 do not.

### gcPCA avoids hyperparameters by including a normalization factor

Our goal for gcPCA was to eliminate the need for hyperparameters and provide unique, correct solutions. To mitigate the bias toward high-variance dimensions, we introduce a normalization factor, such as the total variance in both conditions, which can be calculated by summing the covariance matrices (**C**_*A*_ + **C**_*B*_). The objective function then becomes (eq. 8):

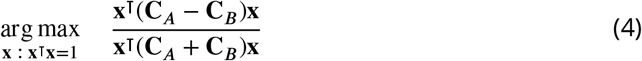

Solving this problem is slightly more complicated than with PCA or cPCA, but ultimately it can also be reduced to a computationally efficient eigenvalue problem (see Methods). The resulting generalized contrastive PCs (gcPCs) maximize relative, rather than absolute, changes in variance between conditions *A* and *B*. This effectively handles the bias toward high-variance dimensions and successfully extracts the ground truth dimensions in our synthetic data, even with finite sampling (Fig. 1E, G), and even when the range of acceptable *α* is narrow (Fig. S1).

This creates two minor complications to be aware of: first, unlike PCA or cPCA, gcPCs are not orthogonal by default. If orthogonality is important for a particular application, we have implemented versions of gcPCA with an orthogonality constraint (see Methods). Second, the normalization factor can create numerical instability if the data are rank-deficient, yielding dimensions with zero or near-zero variance in the denominator. Our implementation of gcPCA prevents this by detecting and excluding those dimensions before performing the calculation (see Methods).

### The open-source gcPCA toolbox contains multiple gcPCA variants enabling optimal handling of diverse use cases

We developed an open-source gcPCA toolbox with implementations in Python and MATLAB of several different variants of gcPCA. This toolbox is freely available at: https://github.com/SjulsonLab/generalized_contrastive_PCA. Here we will present the different variants of gcPCA and their use cases.

#### gcPCA v1.0: traditional cPCA

For version 1.0, we include an implementation of the original cPCA algorithm that finds cPCs maximizing the objective function in Eqn. 3.

#### gcPCA v2.0: gcPCA maximizing A/B

Here we include an implementation that finds gcPCs maximizing the ratio of variance in A to B:

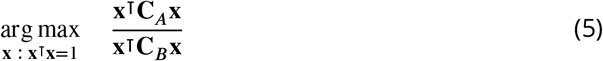

Like cPCA, this method is asymmetrical, meaning it is suitable for situations in which *A* is a foreground condition and *B* is a background condition; in other words, *A* is presumed to be equal to *B* with a low-dimensional pattern added, and the goal is to extract that pattern. The resulting eigenvalues are the ratio of the variance a given gcPC accounts for in *A* to the variance it accounts for in *B*. Thus, they fall in the range [0, ∞), with gcPCs enriched in *A* having eigenvalues > 1.

#### gcPCA v3.0: gcPCA maximizing (A-B)/B

The second method developed is also asymmetrical but based on a relative change:

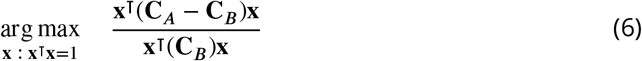

This method is closely-related to v2.0 and is suitable for scenarios in which the investigator wishes to define the gcPCs based on a relative change to a background condition (i.e., finding a 30% increase in the neural activity in condition *A* relative to *B*). The eigenvalues returned are in the range [−1, ∞), with gcPCs enriched in *A* having eigenvalues > 0.

#### gcPCA v4.0: gcPCA maximizing (A-B)/(A+B)

The last of the three methods is based on a relative change:

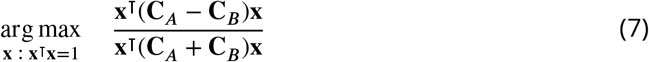

This method is symmetrical, treating conditions *A* and *B* equally, and is appropriate for contrasting conditions in which *B* is a distinct condition and not merely a background to be removed, for example comparing neural data in sleep vs. wakefulness. The eigenvalues are in the range [−1, 1] and are easily interpretable as a traditional index of the form (*A* − *B*)/(*A* + *B*): 1 means that a gcPC only accounts for variance in *A*, -1 means it only accounts for variance in *B*, and 0 means it accounts for equal variance in both. This method is fully symmetrical in the sense that switching *A* and *B* will yield the same gcPCs with the signs of the eigenvalues reversed.

#### gcPCA v2.1, v3.1, and v4.1: Orthogonal gcPCA

Unlike PCs or cPCs, gcPCs are not orthogonal by default. Because orthogonality may be important for some applications, we also include versions of gcPCA with an orthogonality constraint (see Methods). gcPCA v2.1 is the orthogonal version of 2.0, and v3.1 is the orthogonal version of v3.0, and v4.1 is the orthogonal version of v4.0.

#### Sparse vs. dense gcPCA

In high-dimensional datasets, it is often desirable to perform feature selection for easy interpretation of the results. With that in mind, we have also developed sparse versions of gcPCA. This method works by first finding the gcPCs based on the objective function selected, then performing feature selection using an *L*1 lasso penalty (see Methods). All non-orthogonal versions of gcPCA can be run in either sparse or dense mode, but sparsification cannot be used with the orthogonality constraint (***Zou et al., 2006***).

**Table 1.** gcPCA variants in the gcPCA toolbox.

| gcPCA version | Symmetric | Orthogonal | Sparse solution | Note |
| --- | --- | --- | --- | --- |
| **v1** | ✗ | ✓ | ✓ | Equivalent to cPCA |
| **v2** | ✗ | ✗ | ✓ | Objective function 5 |
| **v2.1** | ✗ | ✓ | ✗ | Objective function 5 |
| **v3** | ✗ | ✗ | ✓ | Objective function 6 |
| **v3.1** | ✗ | ✓ | ✗ | Objective function 6 |
| **v4** | ✓ | ✗ | ✓ | Objective function 7 |
| **v4.1** | ✓ | ✓ | ✗ | Objective function 7 |

### Successful extraction of facial expression features with gcPCA

To illustrate the utility of gcPCA with real-world datasets, we first use the Chicago Face Dataset (***Ma et al., 2015***), which contains faces with different facial expressions. Here we used happy and angry expressions as condition *A* and neutral faces as condition *B* (Fig. 2A). This dataset is useful for two reasons: 1) the categorical separation of facial expressions allows an easy evaluation of the contrastive methods, and 2) the dimensions can be visually inspected for features that are being discovered by the method.

**Figure 2.**
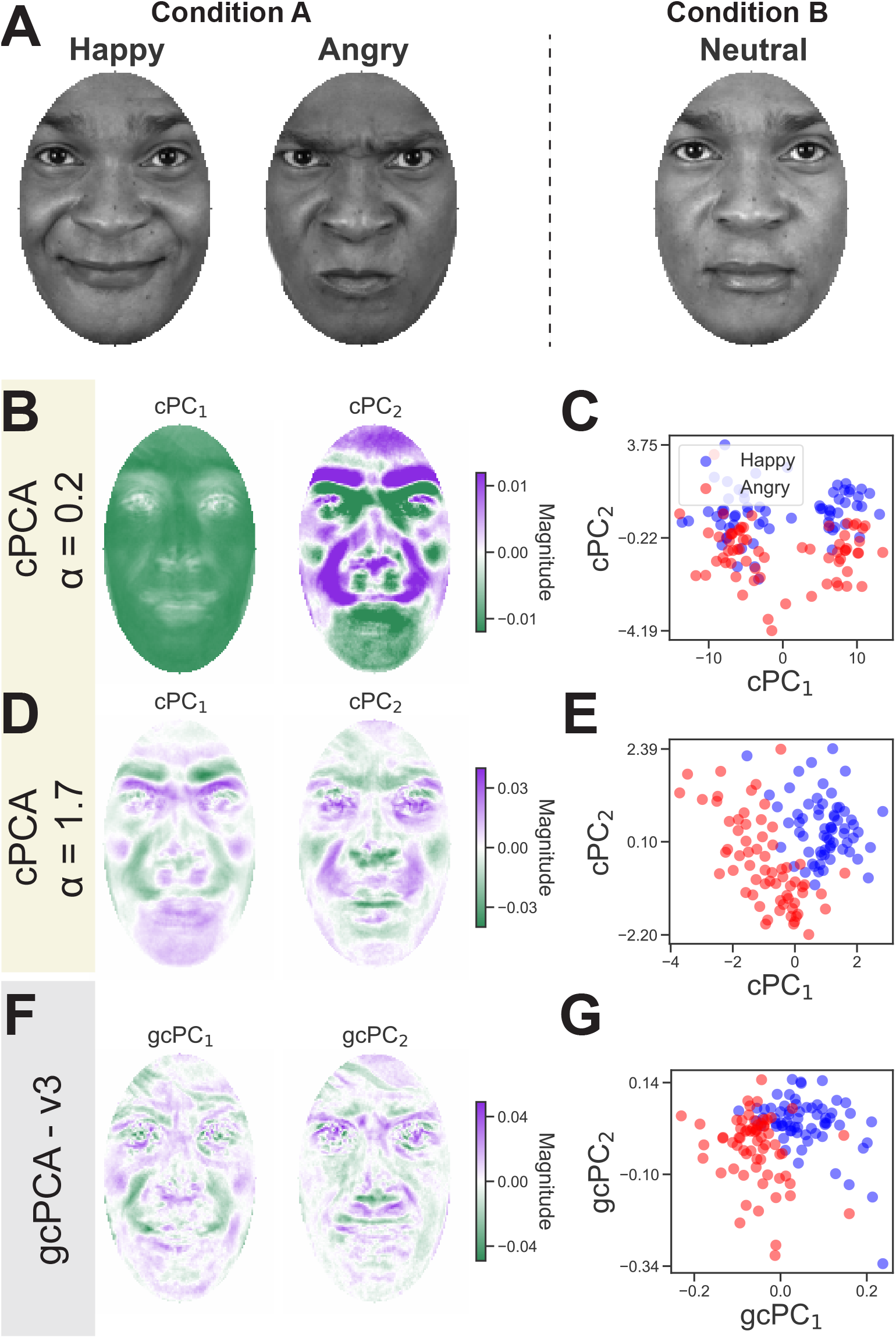
gcPCA correctly extracts contrastive facial features in the Chicago Face Dataset. **A** For this test, contrastive methods were applied to a set of happy and angry face images (Condition *A*) versus neutral face images from the same subjects (Condition *B*). Facial expression changes along the happy-angry axis were therefore the low-dimensional pattern that is enriched in condition *A*. **B** The first *α* identified by cPCA’s algorithm, *α* = 0.2, yields loadings similar to the first PCA dimensions, *i*.*e*. eigenfaces. **C** Projecting the faces onto the cPCs reveals clusters unrelated to the happy and angry facial expressions in condition *A*, indicating an incorrect solution. **D** cPCA with *α* = 1.7 correctly reveals features associated with the expected facial expressions in condition *A*, such as furrowed eyebrows and the region around the mouth and nose. **E** Projecting the faces onto these cPCs reveals the separation of happy and angry faces along the first cPC. Importantly, it was only possible to determine this answer was correct because we knew the labels in advance. **F** Dimensions identified by gcPCA correctly reflect features related to the facial expressions in condition *A*. **G** Projecting the faces onto the first two gcPCs also reveals the separation of happy and angry faces along the first gcPC.

We first applied cPCA to the two conditions and used its automatic *α* selection algorithm to pick two different *α* values. The automatic *α* selection algorithm developed by ***Abid et al. (2018)*** finds representative *α*’s that yield different cPC embeddings so that the investigator can choose the appropriate *α* value. The algorithm is explained fully in the original paper (see supplementary methods - algorithm 2 ***Abid et al. (2018)***). Briefly, it calculates cPCA for an array of different *α*’s (default: 40 different *α* values ranging from 0.01 to 1000 spaced on a log scale), defining a subspace with the top *k* cPCs (default: 2 dimensions), and calculating an affinity matrix between subspaces of different *α*’s. This affinity matrix is then clustered to find *p* clusters, and the medoid of each cluster is a candidate *α*.

The first value returned is *α* = 0.2, and we can see that the first two cPC loadings resemble “eigenfaces” (***Turk and Pentland, 1991***), the largest principal components of facial images (Fig. 2B). This suggests that this *α* is too small, leading cPCA to extract the highest-variance components in condition *A*. Looking at the individual faces projected on these cPCs, two clusters can be identified that separate cleanly along cPC1 (Fig. 2C). However, this solution is incorrect because these clusters do not reflect the two facial expressions comprising condition *A*. Instead, they represent skin color, which should account for equal variance in both datasets because images of the same subjects are present in both conditions. For the second *α* value returned (*α* = 1.7), the cPC loadings exhibit features specific to condition *A* (Fig. 2D), and data projected onto these cPCs recovers the different expressions (Fig. 2E). It is important to note that if we did not have the class labels *a priori*, we could easily believe *α* = 0.2 was the correct answer because it produces better clustering than for *α* = 1.7 (Fig. 2C,E). Using gcPCA v3.0 (asymmetric, non-orthogonal) also reveals features specific to condition *A* (Fig. 2F), and the two expressions in the dataset can be distinguished by their projection onto gcPC_1_ (Fig. 2G), without the requirement of fine-tuning a hyperparameter.

### Applying gcPCA to neurophysiological recordings reveals hippocampal replay without *a priori* knowledge of replay content

A key application for gcPCA is neuronal recordings, which frequently contain hundreds of isolated single units (***de Oliveira et al., 2022; Jun et al., 2017***). A well-studied neurophysiological phenomenon is hippocampal replay, in which hippocampal neurons encoding spatial trajectories traversed during a behavioral task “replay” the same activity patterns in post-task periods (***Wilson and McNaughton, 1994***). It is important to note that spatial location is a continuous variable, so we are not expecting gcPCA or cPCA to find dimensions that cluster the activity into different groups, as with the facial expression data. Instead, we are hoping to see replay of the firing patterns that encode the linear track the animal recently explored (***Wilson and McNaughton, 1994; Foster, 2017***). For our analysis, we used a previously published dataset recorded from hippocampal CA1 ***Girardeau et al. (2017)*** where rats learn the location of an aversive air puff on a linear track. The air puff is only delivered when the rat is running in one direction, called the danger run, and the other direction is the safe run (Fig. 3A). Using previously-established methods (***Kudrimoti et al., 1999***), ***Girardeau et al. (2017)*** found that hippocampal neurons exhibit reactivation of the task representation in post-task activity when compared to pre-task activity. We tested whether cPCA or gcPCA could extract hippocampal replay directly from neuronal activity by contrasting post-task activity (condition *A*) with pre-task activity (condition *B*) (Fig. 3B). For this, we used cPCA and gcPCA to extract cPCs or gcPCs from the pre- and post-task data, then projected the during-task data onto the cPCs/gcPCs to test whether any spatial structure was discernible.

**Figure 3.**
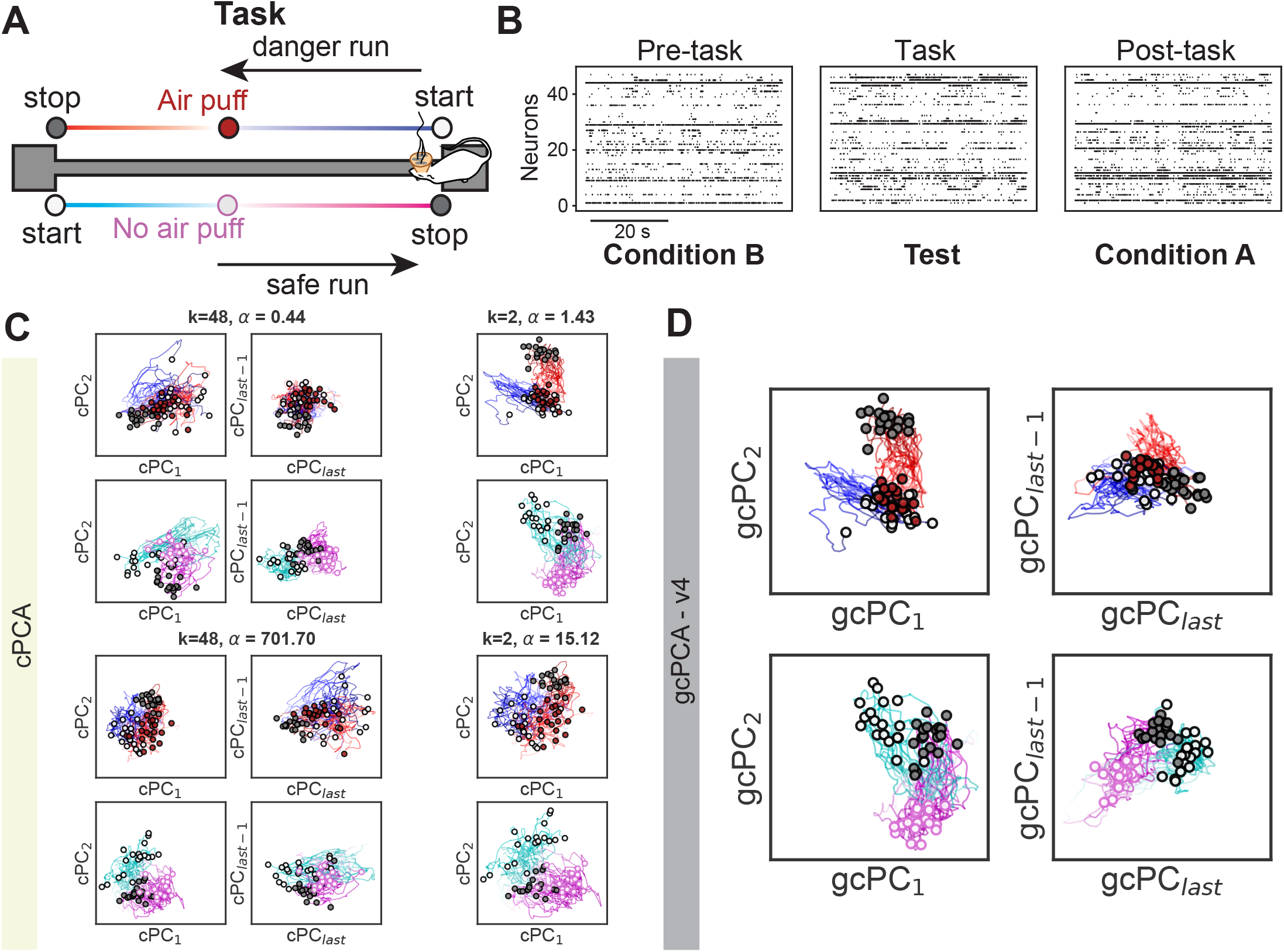
gcPCA v4.0 correctly identifies hippocampal replay in neurophysiological data without prior knowledge of replay content. **A** In ***Girardeau et al. (2017)***, rats were trained to traverse a linear track where one direction has an aversive air puff and the other does not. Rats learned the location of the air puff and which direction was dangerous or safe. **B *Girardeau et al. (2017)*** recorded hippocampal neurons in pre- and post-task periods and used activity recorded during the task as a template to determine that that activity was “replayed” during post-task. We reanalyzed their published data using post-task activity as condition *A* and pre-task activity as condition *B* to identify the dimensions most enriched in post-task activity without taking task-related activity into account. **C** We first performed cPCA, then projected task-related neuronal data onto the cPCs to determine whether task-related data was enriched in the post-task period. The automatic *α* selection algorithm from cPCA returns various *α* values depending on the number of cPCs requested (parameter *k*). *Left Column* Representative *α* values returned with *k*=48 cPCs. cPC_1−2_ are the dimensions most enriched in post-task, and cPC_last_ and cPC_last-1_ are the most enriched in pre-task. No discernible spatial structure is identified, indicating that replay was not detected. *Right Column* With *k*=2, the *α* values returned are of different magnitudes, and one of them (*α* = 1.43) reveals spatial structure related to the task, indicative of hippocampal replay. **D** gcPCA readily identifies the replay of the spatial task structure (left) with no parameter search.

Using cPCA, it was not straightforward to detect replay. When we requested the cPCA algorithm evaluate all the dimensions (*k* = 48), the *α* values identified by the automatic selection did not reveal any obvious task-related spatial structure (Fig. 3C – left column, *α* values 0.44 and 701.70). When we requested a smaller set of dimensions (*k* = 2), the automatic *α* selection returned several different values, with one of them (*α* = 1.43) revealing spatial structure in the task data (Fig. 3C - right column). This reveals that although the only hyperparameter for cPCA is *α* in theory, the automatic alpha selection algorithm depends on *k*, the number of components requested, constituting a second *de facto* hyperparameter.

Applying gcPCA v4 (symmetrical, non-orthogonal), we readily recovered replay of task-related spatial structure (Fig. 3D, top). Importantly, traditional replay analyses require knowledge of the firing patterns during the task to test whether they were overrepresented in post-task sleep (***Kudrimoti et al., 1999; Wilson and McNaughton, 1994; Foster, 2017***). gcPCA was able to extract signatures of replay without prior knowledge of the task-related activity. This may prove useful in many analogous situations in which the experimenter does not have prior knowledge of the pattern they are searching for.

We also took advantage of the symmetric nature of gcPCA and investigated the patterns enriched in pre-task activity, gcPC_last_ and gcPC_last-1_. As expected, they did not exhibit task-related spatial structure (Fig. 3D, top), suggesting they contain replay of environments other than the linear track.

### Applying gcPCA to scRNA-seq data prioritizes disease genes and reveals disease heterogeneity in type II diabetes

Another key application of gcPCA is high-dimensional omics datasets, which often reflect a comparison of two conditions (*e*.*g*. disease vs. healthy). For this analysis, we used published single-cell RNA sequencing data from pancreatic beta cells taken from healthy controls or patients with type II diabetes (T2D)***Martínez-López et al. (2023)***). We used T2D beta cells as condition *A* and healthy control beta cells as condition *B* to test whether gcPCA 4.0 could identify groups of genes that vary more among T2D patients or controls. gcPC_1_ and gcPC_2_ therefore represent axes along which cells from patients vary more and the gcPC_last_ and gcPC_last-1_ represent axes along which control cells vary more. We found that T2D and control data had similar levels of variability (Fig. 4A), but in T2D patients this variability exhibited clear donor-based clustering (Fig. 4A, top left) that was not observed in controls (Fig. 4A, bottom right). Several of the genes with the highest loadings on gcPC_1_ have been previously implicated in T2D (*IMMP2L* (***Diabetes Genetics Initiative of Broad Institute of Harvard and MIT, Lund University, and Novartis Institutes of BioMedical Research et al., 2007; Greenwald et al., 2019***), *STMN1* (***Horn et al., 2016***), *PDHA1* (***Srinivasan et al., 2010***), *CLU* (***Kim et al., 2001, 2006***), *DDIT3* (***Li et al., 2022; Yong et al., 2021***), *SSTR5-AS1*, (***Jian and Felsenfeld, 2018***), *TFF3* (***Fueger et al., 2008***), Fig. 4B, red), notably the top two hits, which were the *TMEM176A/B* genes that ***Martínez-López et al. (2023)*** demonstrated are functionally important for T2D-related beta-cell function. The fact that gcPCA identifies known T2D-related genes and that clustering is specific to cases suggests that gcPCA is revealing disease heterogeneity (***Ahlqvist et al., 2020***).

**Figure 4.**
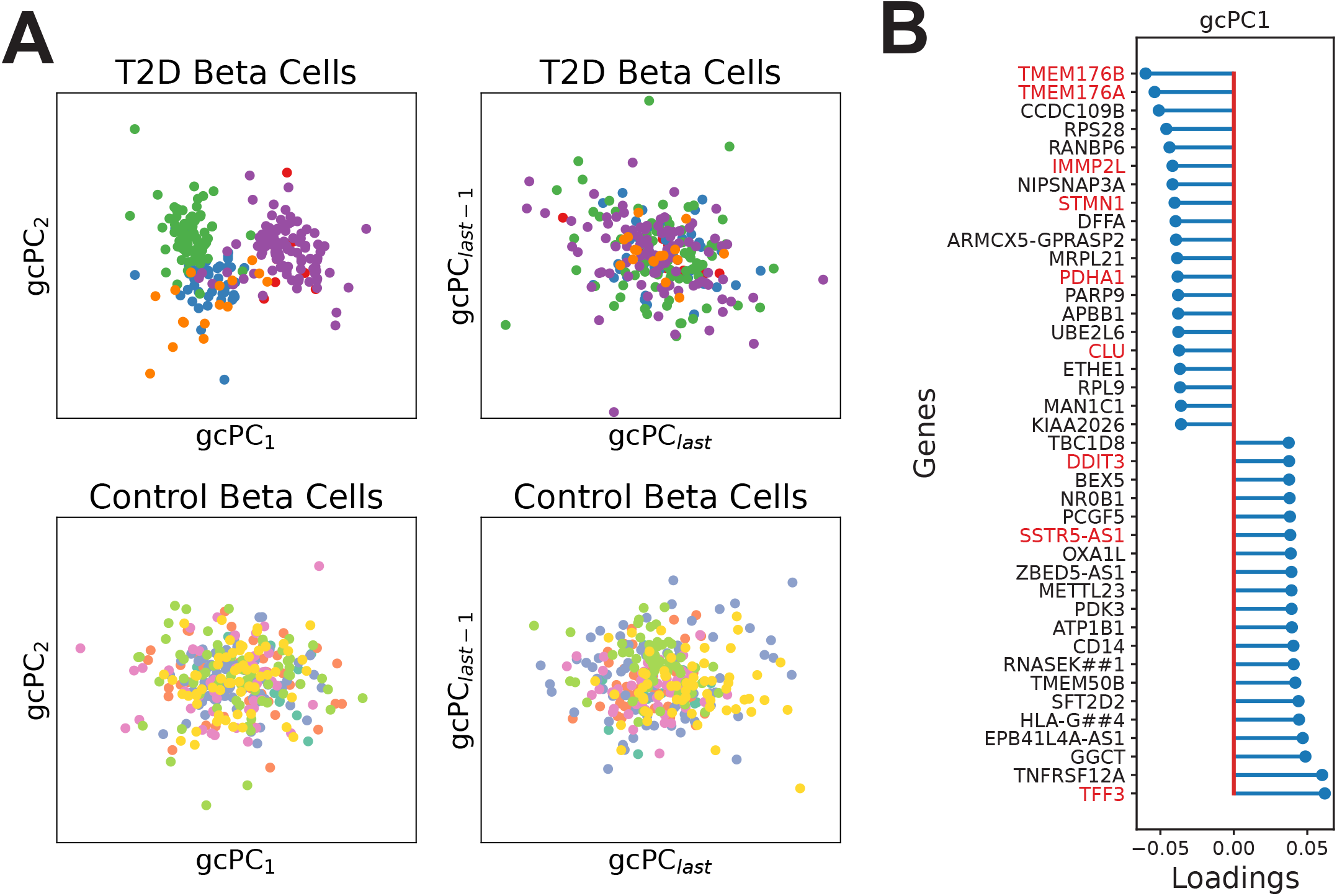
gcPCA v4.0 reveals possible disease heterogeneity in type II diabetes. **A *Martínez-López et al. (2023)*** performed scRNA-seq on isolated pancreatic cells from type II diabetes (T2D) patients and healthy controls. We used gcPCA v4.0 to compare the two conditions and found clustering of beta cells by donor identity (each color is a donor) in gcPC_1,2_ (top left panel). Such clustering was not found in the last gcPCs (top right panel) or in the control donors (bottom left and right panels) **B** The list of 40 genes with the highest loadings on gcPC_1_ includes several previosly linked to T2D (red). Notably, the top two hits, *TMEM176A* and *TMEM176B*, were shown by ***Martínez-López et al. (2023)*** to play functional roles in beta cell function.

## Discussion

Discovering low-dimensional patterns that vary between conditions in high-dimensional datasets is a crucial analysis in many research contexts. Here we present gcPCA, a method that achieves this by examining the covariance structure of the datasets to find dimensions that exhibit the largest relative changes in variance between conditions. This work builds on the pioneering insights in the development of cPCA ***Abid et al. (2018)*** but solves cPCA’s key problem, the requirement for the hyperparameter *α*. Here we showed that the function of *α* is to compensate for bias toward high-variance dimensions in noisy, finitely-sampled data. Further, we showed how this can be circumvented by introducing a normalization factor. Previous work ***Abid et al. (2018)***; ***Boileau et al. (2020)*** has focused on developing and improving methods to find appropriate choices for *α*, but with gcPCA we chose instead to eliminate the *α* hyperparameter entirely. Importantly, the advantage of our approach is not merely that it is computationally cheaper than scanning a range of *α*’s; it is that in most real-world cases there is no way to know whether a given choice of *α* yields a correct solution.

We wish to address a common point of confusion by reiterating how gcPCA differs from LDA or PLS. These are methods that find patterns optimally distinguishing two datasets, but gcPCA finds patterns that exhibit more within-dataset variability in one dataset than another. As a fictitious example, LDA might find a height/weight dimension distinguishing a university rugby team from the general population because rugby players are taller and heavier. In contrast, gcPCA would be more likely to find an age/education-level dimension because those features exhibit more variability in the general population than in a university team. Despite the simplicity of this analysis, it reveals interesting phenomena in high-dimensional biological data such as hippocampal replay (Fig. 3) or transcriptomic heterogeneity in disease states (Fig. 4).

gcPCA has a few caveats: first, unlike ordinary PCs or cPCs, gcPCs are not orthogonal by default. Our toolbox includes versions of gcPCA with an orthogonality constraint (v2.1, v3.1, and v4.1), which comes at increased computational cost because a new eigendecomposition must be performed for each gcPC. Second, the normalization factor introduces the possibility of numerical instability if the denominator matrix is rank-deficient, meaning it has dimensions with zero (or near-zero) variance that create a “divide-by-zero” situation. However, the implementation of gcPCA in the toolbox automatically excludes these dimensions if they exist. Finally, there may be situations in which cPCA’s *α* could be a feature, rather than a bug, if the investigator has prior knowledge that the patterns of interest will lie in high- or low-variance dimensions. Choosing an appropriate *α* could then intentionally bias the analysis in favor of the results of interest. In such cases, it would be relatively straightforward to extend gcPCA by adding a parameter that accomplishes a similar result by adjusting the eigenspectrum of the denominator matrix in the objective function, *e*.*g*. (**C**_*A*_ − **C**_*B*_). There are also other extensions that could be added, such as contrasting more than two conditions or incorporating nonlinearity, which can be relevant to specific data problems. However, we leave the development of such tools for future efforts.

The biological sciences are currently undergoing an explosion of technologies that produce high-dimensional datasets, including novel forms of microscopy and neuroimaging, high-speed video tracking, Neuropixels recordings, -omics approaches with single-cell resolution, and many others. In addition to the analyses of electrophysiological recordings or single-cell RNA sequencing data demonstrated here, gcPCA could be applied to any of these experimental modalities. We thus anticipate that the open-source gcPCA toolbox will provide a valuable resource facilitating a broad range of biological investigations that require contrasting two experimental conditions.

## Methods

### Generalized contrastive PCA

Our motivation for the following method stems from eliminating the necessity of the free parameter *α* in the contrastive PCA method. To accomplish this, we introduce a normalization factor to mitigate the bias toward high-variance dimensions. We will summarize the process of calculating the gcPCs using gcPCA v4.0 as an example, but v2 and v3 are analogous. gcPCA v4.0 has the following objective function, as shown in equation (eq. 8):

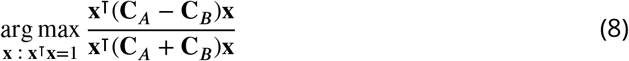

A potential problem of this objective function is the denominator creating numerical instability if there are vectors that have eigenvalues approaching zero in the denominator covariance matrix. To address this, we consider only the principal components (*j*) of that matrix that have non-zero eigenvalues. In this case, the matrix *j* is composed of the principal components of the row-wise concatenated datasets *A* and *B* that have non-zero eigenvalues. We then substitute for **x** using **x** = **J***γ*, yielding:

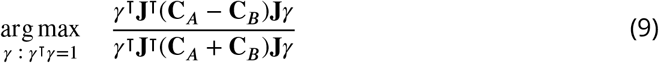

The matrix (**J**^⊺^(**C**_*A*_ + **C**_*B*_)**J**) in the denominator is guaranteed to be positive definite, allowing us to find a symmetric matrix **M** that is its square root, yielding equation (eq. 10):

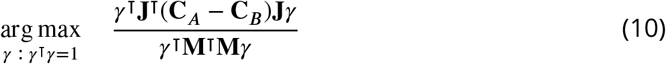

Let **y** = **M***γ*, yielding:

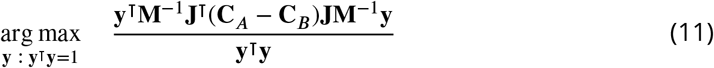

This optimization problem can be solved with the eigendecomposition of the numerator matrix. The vectors in **X** are then calculated using equation (eq. 12):

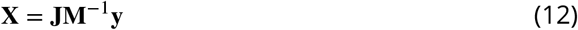

The column vectors in **X** are referred to as gcPCs, following the term cPCs used in ***Abid et al. (2018)***. The other versions of gcPCA can be solved in the exact same way, by replacing **C**_*A*_ − **C**_*B*_ with **C**_*A*_ (v2) and replacing **C**_*A*_ + **C**_*B*_ with **C**_*B*_ (v2 and v3).

In our case, the gcPCs (**X**) are not guaranteed to be orthogonal due to the presence of the **M**^−1^ matrix. By default, gcPCA returns non-orthogonal gcPCs. If orthogonality is desired (as in gcPCA v4.1), we iteratively shrink matrix **J** to remove the subspace spanned by the gcPCs (*i*.*e*. columns of **X**) that have already been found. At each step, we compute the largest eigenvector **x** of equation 11, then project it into the feature space with equation 12 and concatenate it column-wise into the growing matrix **X**. To shrink **J**, we first regress out the **x** from **J**, as shown in equation 13:

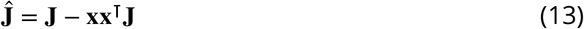

We then use SVD to get the left singular vectors of Ĵ, and we define 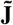 as the first *n* − *i* of these (on the *i*-th iteration). 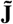 serves as an orthonormal basis for the subspace of **J** that is orthogonal to **X**, and we use 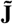 as the new **J** in eq. 9 for the next iteration. This process continues until *n* gcPCs are found, which can be the number of features in the dataset, the minimum rank of the conditions, or a number specified by the user for the gcPCs to be extracted.

### Sparse gcPCA

We developed an extension for sparse gcPCA using a similar approach as sparse PCA (***Zou et al., 2006***) and sparse cPCA (***Boileau et al., 2020***). Here we will first review the sparse PCA framework, then explain how we adapt it for gcPCA.

#### Sparse PCA method

Sparse PCA was first proposed as a reinterpretation of PCA as a regression problem. In brief, given a matrix **X**_*n*×*k*_, where the first *k* ordinary PCs are organized column-wise and are orthonormal, PCA can be seen as minimizing the following objective:

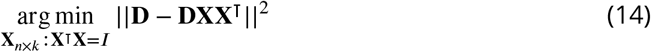

Where **D** is the data matrix of size features × samples. To achieve sparse loadings in the first *k* PCs, ***Zou et al. (2006)*** proposes the use of elastic net regularization, as shown in the following objective function:

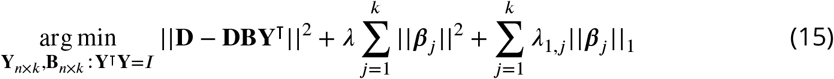

Where **B** is the matrix containing the sparse PCs *β*_*j*_, and **Y** is a column-wise matrix that projects data from the sparse PC space to the feature space. The elastic net is the combination of the ridge penalty 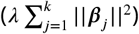 and the lasso penalty 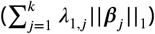. The ridge penalty is used to correct a rank-deficient matrix **D** for numerical purposes. In ***Zou et al. (2006)***, the same λ was used for all *k* components while using a different λ_1,*j*_ for every component. To numerically solve equation 15, (***Zou et al., 2006***) use an alternating algorithm in which **Y** is held constant as we solve for **B**, then **B** is held constant while we update **Y**, and this is repeated until the algorithm converges. **Y** is initially set to be equal to the ordinary PCs (**X**), then we can find each column of **B** by the following elastic net regression:

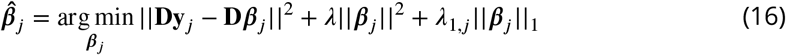

Next, **B** is fixed, meaning the penalty terms can be ignored, and the new **Y** is defined as:

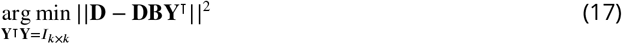

The solution to 17 can be found by a reduced rank form of Procrustes rotation. Using SVD, we find

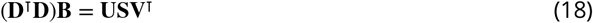

We then set Ŷ= **UV**^⊺^. To solve eq. 16, ***Zou et al. (2006)*** has shown that is only necessary to know the Gram matrix **D**^⊺^**D**. For a fixed **Y**, finding *β*_*j*_ is equivalent to minimizing:

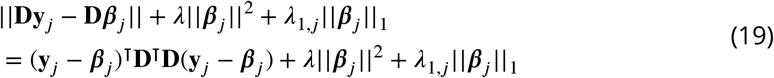

If the covariance matrix of **D** is known (denoted below as **Σ**), the term **D**^⊺^**D** can be replaced with **Σ**. For solving the eq. 16, the **D** matrix can be replaced by 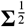, which is the square root matrix of **Σ**. Resulting in the updated equation 20

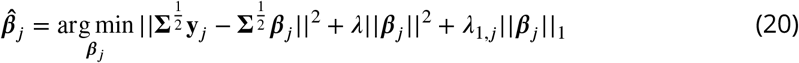

#### Sparse gcPCA method

We can implement sparse gcPCA by adapting the sparse PCA method presented and following similar steps proposed in ***Boileau et al. (2020)***. For sparse gcPCA, the covariance matrix **Σ** is replaced with the matrix **Θ** which reflects the appropriate objective function of the version used. Following gcPCA objective function 11, **Θ** = **M**^−1^ (**C**_*A*_ − **C**_*B*_) **M**^−1^, where **M** is the square root matrix of **C**_*A*_ + **C**_*B*_. For version 1 (equivalent to cPCA), we instead use **Θ** = **C**_*A*_ − *α***C**_*B*_, and for versions 2 and 3 we change **Θ** to match their respective objective functions, as mentioned previously. We removed the **J** matrix from the objective function so the sparsity is enforced in the features and not in the principal components. The components **X** are the gcPCs identified by the ordinary gcPCA algorithm. Following the numerical solution for sparse PCA presented before, the sparse gcPCA is obtained by the following alternating algorithm until convergence:

**B given Y:** Each sparse gcPC (denoted here as *β*_*j*_) was found according to the following elastic net solution:

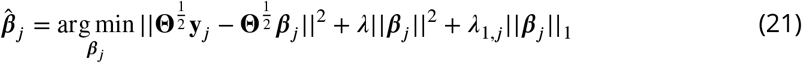

Where **y**_*j*_ is the *j*^th^ gcPC, and *β* is the sparse gcPC. The ridge penalty λ is used to fix rank-deficient matrices. To simplify our approach, we used the same λ_1_ for all components instead of a different λ_1,*j*_ for every *j*^th^ component. Therefore, the eq. 21 is reduced to a lasso regression and is solved through least angle regression, similar to ***Zou et al. (2006)***.

**Y given B:** Using fixed **B**, we can find a new Ŷwith a Procrustes rotation using SVD:

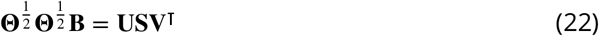

We can then determine Ŷ= **UV**^⊺^. These steps are repeated until loadings converge. The main caveat with this approach is that for gcPCA v3 or v4, the matrix **Θ** can have negative eigenvalues, which prevents the calculation of the square root matrix 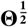. To overcome this problem we follow similar steps to ***Boileau et al. (2020)***, which we briefly replicate here. In gcPCA v3 or v4, positive and negative eigenvalues have a clear interpretation: for the matrix **Θ**, positive eigenvalues denote vectors with larger variance in condition *A*, while negative eigenvalues denote vectors with larger variance in condition *B*. We can then perform eigendecomposition of **Θ** and replace any negative eigenvalue with zeros to make **Θ**_+_ positive semi-definite:

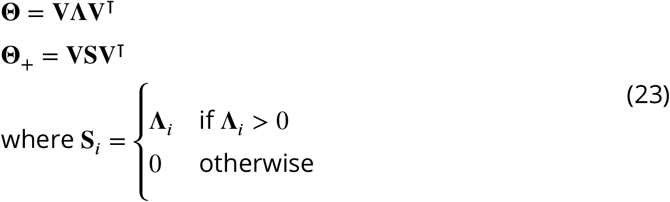

We can then define the square root matrix 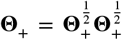. We note that this matrix is only able to solve sparse gcPCA for vectors enriched in condition *A*. Here we propose a solution for finding the vectors also in condition *B*. As mentioned previously, the sign of the eigenvalues of matrix **Θ** indicates whether they are more expressed in condition *A*(+) or *B*(-). To solve the sparse gcPCA for condition *B*, we turn any positive eigenvalue to zero and switch the sign of negative eigenvalues to positive:

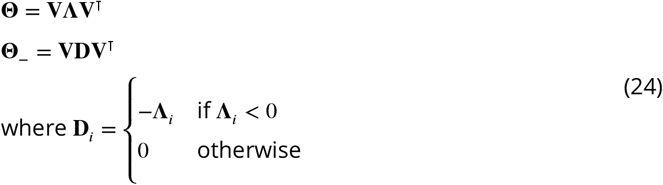

Although **Θ**_−_ does not contain negative eigenvalues, it still represents the dimensions most expressed in condition *B*. This procedure is equivalent to switching the order of conditions *A* and *B* as the eigenvalues would be flipped in sign. The sparse gcPCs are found separately for **Θ**_+_ and **Θ**_−_ and are later concatenated for the final sparse gcPCs.

### Synthetic data generation

In the synthetic data, we generated two conditions with 1 × 10^5^ samples and 100 dimensions. The dimensions were sampled from a Gaussian distribution (mu = 0 and sigma = 1) and then orthogonalized using singular value decomposition and picking the left singular vectors. In each condition, we created a pattern in the samples that was to be discovered. In condition *A*, we took dimensions 71 and 72 and drew the samples from a uniform distribution ([0, 1]). In dimension 71 we replaced any value from 0.3 and 0.7 with a different uniform distribution ([0, 0.4]). In dimension 72 all the values between 0.4 and 0.6 were replaced with another uniform distribution ([0, 0.4]). This created the square with a square hole in the middle in Fig. 1 A. The values were then offset by 0.5, the samples were sorted by the angle they formed in each dimension, calculated by the inverse tangent (tan inverse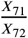). The samples in each dimension were normalized by their *l* -norm. In condition *B*, we generated the samples of dimensions 81 and 82 from a uniform distribution ([0, 1]), sorted the sample values based on dimension 81, and rotated both dimensions by 45 degrees. This created the diamond shape seen in Fig. 1 A. The sample values were later normalized by their *l*_2_-norm. The samples for all the other dimensions were drawn from a Gaussian distribution (*μ* = 0 and σ = 1), and then normalized by their *l*_2_-norm. The magnitude of each dimension was established as a line with a negative slope, starting at value 10 in the 1st dimension and ending at 0.001 in the 100th dimension. For condition *A*, we doubled the magnitude in dimensions 71 and 72, while in condition *B* we doubled the magnitude in dimensions 81 and 82. Identifying these changes in magnitude is the goal of contrastive methods. Because the samples were drawn from a normal distribution, the dimensions will display correlations among them. To estimate the total variance explained by each dimension, we use a QR decomposition approach described in ***Zou et al. (2006)***. In brief, let *Z* be a matrix containing scores of each dimension generated, the variance is usually calculated through tr(*Z*^⊺^*Z*), where tr is the trace of the matrix. However, in correlated scores, this estimate is too optimistic. Using regression projection, it is possible to find the linear relationships of the dimensions and correct to find the adjusted total variance. ***Zou et al. (2006)*** shows that this is equivalent to using QR decomposition in *Z*, such that *Z* = *QR* where *Q* is orthonormal and *R* is upper triangular, and calculating the adjusted variance as follows:

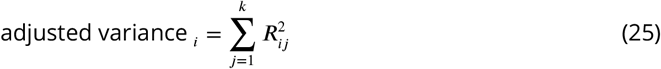

### Face dataset

For the facial expression analysis, we used the Chicago Face Database ***Ma et al. (2015)***. This database consists of neutral and emotional expression faces, and for the analysis we used a subset of samples that had happy and angry faces alongside neutral ones. We used only male faces to reduce variability in feature positioning. The images used were cropped by an ellipse (length of 75 pixels and width of 45 pixels) centered in the face to focus on the facial expression rather than other features such as hairstyle or shoulders. Each sample image was flattened from a two-dimensional matrix to a vector, and all the flattened samples were then concatenated, resulting in a matrix of samples x features. Each feature was z-scored and normalized by its *l*^2^-norm. Condition *A* consisted of all the samples of happy and angry facial expressions, while condition *B* samples were neutral expressions.

### Hippocampal electrophysiology data

We used a previously published hippocampal electrophysiology dataset, with the experimental details listed in the original publication ***Girardeau et al. (2017)***. In brief, Long-Evans rats were implanted with silicon probes in the dorsal hippocampus CA1 region (either left or right hemisphere), and neuronal activity was isolated through automatic spike sorting and manually curated. Animals were trained to collect water rewards at the end of a linear track, and an air puff was introduced at a fixed location for every lap, in only one of the directions. Recordings consisted of task, where the animal learned the air puff location, and periods of pre- and post-task activity. For testing the contrastive methods, we used pre-task recordings as condition *B* and post-task recordings as condition *A*. For our analysis, we only used neurons that had a minimum firing rate of 0.01 spikes/s during the task. We binned the neural data using a bin size of 10 ms and smoothed using a rolling average with a Gaussian window of size 5 bins. The data was then z-scored and normalized by the norm before testing the contrastive methods. The task data was then projected on the contrastive dimensions for evaluation.

### Pancreatic single-cell RNA sequencing

For the single-cell RNA sequencing data analysis, we used a previously published dataset ***Martínez-López et al. (2023)***, available at GEO accession GSE153855, consisting of scRNA-seq data from human pancreatic islet cells from patients with type II diabetes and healthy controls. We used the annotated dataset to identify the beta cells, which were identified previously by the authors ***Martínez-López et al. (2023)***. For condition *A* we used the beta cells from subjects that had type II diabetes, and for condition *B* we used the beta cells from healthy patients. We used the expression values in reads per kilobase of the gene model and million mappable reads (RPKMs). The values were log-transformed, and all the features were centered before the analysis. gcPCA was performed using the same set of genes used in the analysis by ***Martínez-López et al. (2023)***.

## Acknowledgments

We would like to thank Soyoun Kim, Ruben Coen-Cagli, Ehsan Sabri, Wenzhu Mowrey, Cleiton Lopes-Aguiar, and members of the Sjulson and Batista-Brito lab for valuable conversations and insightful comments on the manuscript. This work was supported by funds from the National Institute on Drug Abuse (DP1 DA051608 and R01 DA051652 to LS), as well as from the Whitehall Foundation and McManus Charitable Trust.

## Author contributions

L.S. and E.F.O. conceived the project and developed the mathematical framework. E.F.O., P.G., and L.S. wrote the toolbox code and performed data analysis. L.S. supervised the project with assistance from J.H.L. and R.B.B.

---

**Figure 1–figure supplement 1.**
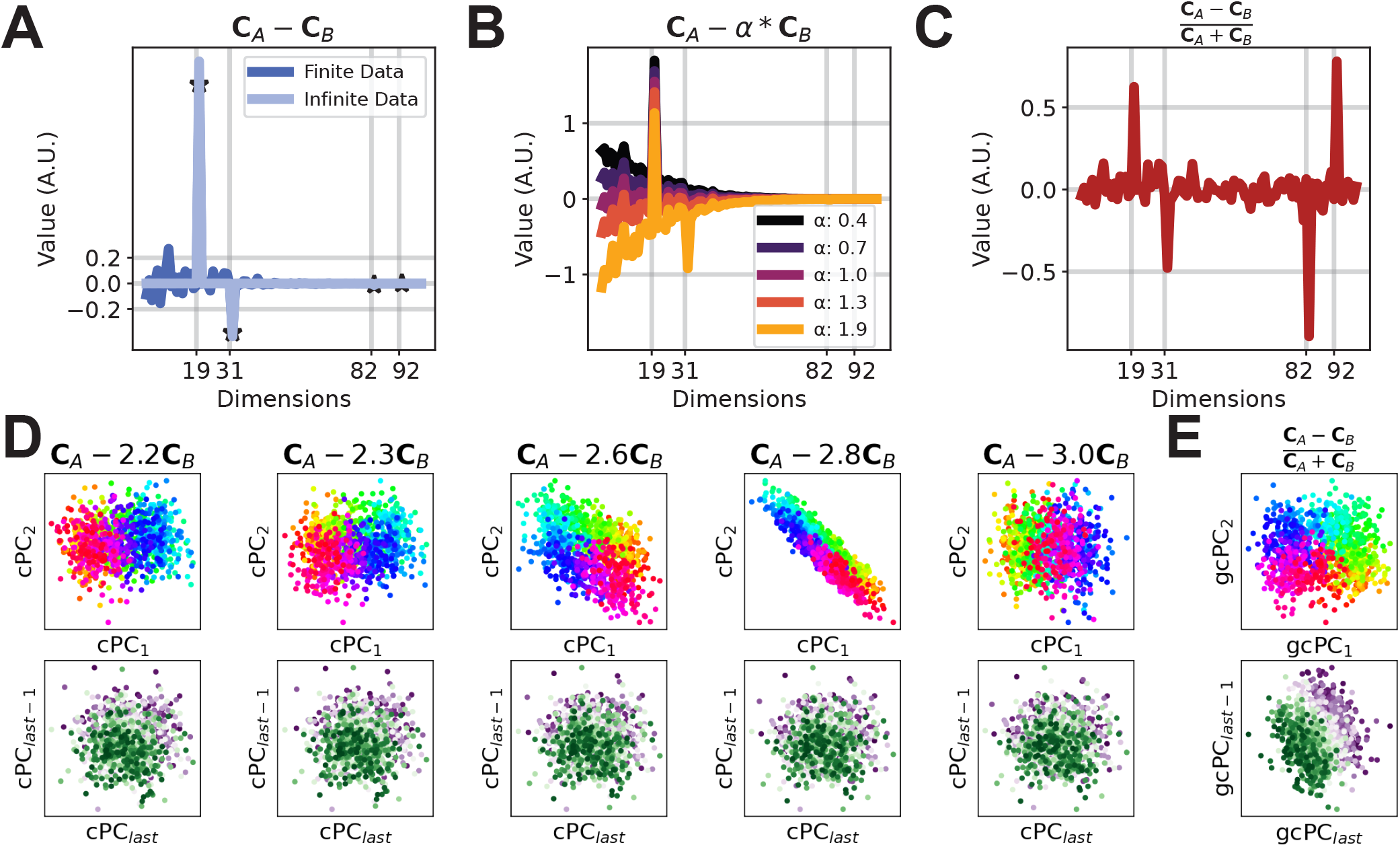
The range of cPCA *α* values yielding correct solutions can be very narrow. **A** We generated synthetic data with enriched variance in both high and low variance dimensions simultaneously. In condition A we enriched the variance of two dimensions (dimensions 19 and 92), and condition B in two other dimensions with high and low variance (dimensions 31 and 82). This panel shows the finite and infinite data results for **C**_*A*_ − **C**_*B*_. Stars represents the finite data value in the enriched dimensions. Even though the high variance dimensions are easy to detect with this method, the low variance ones are still occluded by spurious variability in high variance dimensions. **B** cPCA can reveal enriched high variance dimensions, but enriched low variance dimensions are hard to identify. **C** gcPCA can find all enriched dimensions simultaneously for conditions *A* and *B*. **D** The range of *α* values yielding the correct solution becomes narrow because the enriched dimensions have different absolute variance. **E** gcPCA correctly identifies all enriched dimensions in both conditions.

## Notes

### Competing Interest Statement

The authors have declared no competing interest.

